# Extending the MARTINI 3 Coarse-Grained Forcefield to Polypeptoids

**DOI:** 10.64898/2026.04.10.717689

**Authors:** Jiaxin Wang, Zhou Yu, Mingfei Zhao

**Affiliations:** Department of Chemical and Biological Engineering, The University of Alabama, Tuscaloosa AL USA, 35487; Department of Mechanical Engineering, The University of Alabama, Tuscaloosa AL USA, 35487

## Abstract

Polypeptoids (poly-N-substituted glycines) are synthetic peptidomimetic polymers with their sidechains attached to the backbone amide nitrogen rather than the α-carbon in natural peptides. Peptoids display pronounced sequence-dependent conformational flexibility arising from the absence of backbone hydrogen bonding and slow cis/trans ω-dihedral isomerization. Despite growing interest in peptoid-based biomaterials, a coarse-grained (CG) model compatible with the modern MARTINI 3 framework is not yet available, limiting mesoscale simulation of peptoid structure and self-assembly. In this work, we develop the first MARTINI 3 compatible peptoid CG forcefield, covering 19 commonly used residue types. Extensive all-atom reference simulations employing parallel bias metadynamics (PBMetaD) were performed to ensure converged sampling of ω-dihedral transitions. Bonded parameters were derived from atomistic distribution functions via direct Boltzmann inversion (DBI), while nonbonded interactions were primarily adopted from the standard MARTINI 3 parameter library. The resulting CG model reproduces structural and thermodynamic properties in close agreement with all-atom simulations, while providing up to 57-fold enhanced computational efficiency. To facilitate its adoption by the research community, we have integrated all parameters and workflows to the MARTINI-based martinize2 tool, enabling automated generation of MARTINI 3 peptoid structures and topologies. This work establishes a transferable and computational efficient framework for simulating large-scale peptoid confirmations, assemblies, membrane interactions, and nanostructure formation, and supports the rational design of next-generation sequence-specific functional peptoid-based materials.

## 1. Introduction

Polypeptoids, or poly-*N*-substituted glycines, are synthetic biomimetic sequence-defined polymers. Chemically analogous to peptides, peptoids differ in that their sidechains are attached to the backbone nitrogen rather than the α-carbon, which imparts distinct structural and physicochemical properties.^1–3^ This key modification eliminates inter- and intrabackbone hydrogen bonding, giving rise to a family of non-natural polymers whose conformations and properties are entirely determined by sidechain chemistry.^4^ Building upon this, the solid-phase submonomer synthesis method enables efficient construction of peptoids with diverse side-chain chemistries and precisely tunable sequences.^1, 5, 6^ Compared with peptides, peptoids exhibit greater conformational flexibility, enhanced proteolytic stability, excellent thermal processability, and good backbone solubility, as well as unique folding behaviors.^7, 8^ These features have led to broad applications of peptoids in biomedicine and materials science, including their use in drug delivery systems, advanced materials, and the self-assembly of biomimetic nanostructures.^9–13^

Reliable computational prediction of sequence-dependent peptoid structures and dynamics underpins the rational design, optimization, and engineering of peptoid-based materials, with applications spanning drug delivery to enzyme design.^14–16^ Thus far, residue-specific all-atom parameters for diverse peptoid monomers have been derived from common biomolecular force fields such as GAFF, CHARMM, and OPLS.^17–21^ These customized force fields have been employed to predict the free-energy landscapes and stable conformations of peptoid chains and crystals. For example, Berlaga et al. recently developed CHARMM-compatible MoSiC-CGenFF-NTOID, based on CGenFF-NTOID, combining reparametrized backbone conformational parameters with default CGenFF sidechain parameters to modularly extend the force field to all substituted-methyl side chains.^22^

However, despite advances in hardware and algorithms, all-atom (AA) molecular dynamics (MD) simulations remain limited to relatively small systems and to sub-microsecond timescales. Therefore, coarse-grained (CG) MD models have been developed as a lower-cost computational alternative. The method bundles atoms into CG beads, reducing atomic-level detail yet preserving the key physical interactions and thermodynamic characteristics to balance efficiency and accuracy.^23–26^ Typically, in comparison to its all-atom counterpart, the CG forcefield smoothens out the energy landscape, thereby helping to avoid local energy-minimum “traps”, and substantially reducing the number of degrees of freedom, thereby enabling exploration of configurational space over much longer time and length scales.^22, 27–29^ Notably, cis/trans isomerization in peptoids is a defining feature that governs backbone flexibility, secondary structure, and biological activity, enabling them to mimic or modulate protein functions, form stable helical architectures, control folding kinetics, and serve as programmable scaffolds for drug design, in contrast to peptides where the trans form dominates.^30^ Control of this isomerization through sidechain engineering further expands their functional tunability. At the same time, the large activation barriers associated with backbone amide cis/trans interconversion lead to rare-event dynamics that are difficult to converge within feasible all-atom simulation timescales, even with enhanced sampling. By smoothing the torsional energy landscape and extending accessible timescales, CG models provide a practical route to efficiently sample cis/trans transitions and their collective effects on peptoid conformational behavior.

The MARTINI model is among the most popular CG models in biomolecular simulation due to its transferable bead types and modular “building-block” design, enabling applications across lipids, sterols, carbohydrates, nucleic acids, proteins, and small molecules.^31, 32^ The recent release of the MARTINI 3 further advances CG modeling by introducing refined bead sizes, a unified backbone representation, and chemically specific interaction labels, while rebalanced nonbonded interactions significantly reduce the excessive aggregation observed in earlier versions.^33^ In parallel, CG models for polypeptoids have evolved from bespoke representations to more transferable frameworks. Early models such as MFCGTOID successfully capture nanosheet formation but lacked of compatibility with standard CG forcefields.^23, 34, 35^ Subsequent efforts introduced MARTINI-compatible peptoid models, where bottom-up parametrization strategies based on iterative Boltzmann inversion and potential of mean force matching enabled the accurate reproduction of structural and thermodynamic properties.^29, 36–40^ Importantly, enhanced sampling methods were required to capture slow cis/trans isomerization of the peptoid backbone.^29^

More recent efforts have focused on improving transferability and chemical diversity. The Martinoid forcefield combined bottom-up parametrization with top-down refinement against experimental partition coefficients (log *D*7.4), enabling broader coverage of monomer chemistries and sequence-dependent behavior.^41^ Most recently, a generalizable multiscale workflow based on relative entropy minimization has been proposed to refine CG parameters to minimize information loss relative to atomistic reference system without experimental input.^42^ Despite these advances, existing peptoid CG models remain largely confined to MARTINI 2 or bespoke framework, often requiring modified bead definitions or system-specific parameter tuning, which limits transferability and extensibility.

Notably, extension of peptoid CG models to the MARTINI 3 framework has not yet been reported. Because MARTINI 3 introduces a rebalanced interaction scheme and an expanded, more chemically consistent bead set,^33^ parameters developed under MARTINI 2 are not directly transferable without compromising compatibility. This motivates the development of a dedicated MARTINI 3-compatible peptoid CG forcefield. In this work, we developed a bottom-up CG peptoid model encompassing 19 residues using a unified and chemically consistent mapping scheme fully compatible with the MARTINI 3 framework. Extensive enhanced sampling was performed at the all-atom level via parallel-bias metadynamics (PBMetaD) to ensure thorough exploration of slow backbone ω-dihedral cis/trans isomerization as well as other relevant conformational degrees of freedom. Bonded interactions of the CG model were derived through direct Boltzmann inversion (DBI) of the AA distribution functions, while nonbonded parameters were largely inherited from the standard MARTINI parametrization with several targeted refinements to improve transferability. The resulting force field was rigorously validated against a suite of atomistic benchmarks, including radius of gyration (Rg) trends, backbone dihedral free energy analysis, dimerization potential of mean force (PMF), and heterotrimer aggregation, showing consistently strong agreement with AA reference data across structural and thermodynamic observables. The resulting CG models for common peptoid classes deliver nearly a 57-fold increase in computational efficiency over all-atom simulations, enabling direct access to nanomaterial behavior and assembly at otherwise inaccessible length and time scales. Furthermore, the workflow has been fully integrated with Martinize2 through a publicly available GitHub release (https://github.com/MZhaoLab/polypeptoid_cg_model), for automated topology generation and broad use within the MARTINI 3 ecosystem. The subsequent sections are structured as follows: Section 2 describes the computational methodology, Section 3 describes force-field model development, Section 4 describes validation results.

## 2. Methods

### 2.1. AA MD Simulations

We conducted AA MD simulations of peptoids with GROMACS 2024.4.^43–45^ The initial peptoid structure files were generated from the MoSiC-CGenFF-NTOID GitHub repository (https://github.com/UWPRG/mftoid-rev-residues), which provides a modular and extensible CHARMM-compatible model for all-atom simulation of polypeptoids.^22^ To ensure chemical completeness, each single peptoid polymer, with chain lengths n = 4, 6, 8, 10, 15, and 20, was capped with acetyl (Ac) groups at the N-terminus and aminomethyl (NHMe) groups at the C-terminus before being placed in a cubic simulation box ranging in size from 6.0 × 6.0 × 6.0 nm³ to 12.0 × 12.0 × 12.0 nm³ using Packmol.^46, 47^ Approximately 7000 to 55000 water molecules were supplied to reach a density of 1.0 g/cm³, with periodic boundary conditions applied in all directions. For systems carrying net charges, sodium (Na⁺) or chloride (Cl⁻) ions were added as needed to maintain charge neutrality. Peptoids were modeled using the MoSiC-CGenFF-NTOID all-atom force field, while water molecules were represented using the TIP3P model.^22, 48^ Prior to equilibration, steepest descent energy minimization was performed to eliminate high-energy atomic clashes, removing any forces exceeding 1000 kJ/(mol·nm). Initial atomic velocities were assigned according to the Maxwell-Boltzmann distribution at 300 K followed by a 4 ns equilibration in the NPT ensemble at 300 K and 1 bar, employing a velocity rescaling thermostat with a time constant of 0.1 ps, along with a Berendsen barostat characterized by a time constant of 1.0 ps and an isotropic compressibility of 4.5 × 10⁻⁵ bar⁻¹.^49, 50^ Subsequently, 500 ns production simulations were performed in the NPT ensemble under the same temperature and pressure conditions, during which temperature was maintained using the Nosé-Hoover thermostat with a coupling constant of 1.0 ps, and pressure was regulated using the Parrinello-Rahman barostat, also with a 1.0 ps coupling time and the same compressibility value.^51, 52^ The leap-frog algorithm was used to integrate Newton’s equations of motion with a time step of 2 fs. All covalent bonds involving hydrogen atoms were constrained using the LINCS algorithm.^53, 54^ Electrostatic interactions were computed using the particle mesh Ewald (PME) method, with a real-space cutoff of 1.0 nm and a Fourier grid spacing of 0.08 nm.^55^ Lennard-Jones interactions were smoothly shifted to zero between 1.0 nm and the cutoff. To ensure sufficient temporal resolution for post-simulation analysis, trajectory data were recorded every 10 ps.

### 2.2. AA PBMetaD Simulations

PBMetaD is widely recognized for its effectiveness in enhancing the sampling efficiency of polypeptoid chains with realistic lengths, playing a pivotal role in generating converged AA reference trajectories critical for CG model parameterization.^36, 37, 56^ In particular, PBMetaD is well suited for sampling the amide w dihedral angles, which are more flexible in peptoids than in peptides and undergo rare cis/trans isomerization events due to a high activation barrier (∼63 kJ/mol).^57–59^ By applying multiple low-dimensional biases in parallel, PBMetaD efficiently overcomes this barrier, enabling thorough exploration of slow degrees of freedom such as ω dihedrals.^57, 60^ In this study, we applied PBMetaD to each peptoid system by introducing bias potentials on all backbone ω dihedral angles, thereby facilitating thorough exploration of conformational space. We have also tested to apply bias potentials to all backbone dihedral angles (φ, ϕ, ω), yet no significant differences were observed compared to ω only. To ensure thorough exploration of torsional transitions, bias potentials were periodically applied across the full angular domain (x ɛ [−π, π)). The adopted parameters were as follows: a Gaussian deposition stride of 500 ps, an initial height (W₀) of 1.2 kJ/mol, a Gaussian width (σ) of 0.3 radians, and a bias factor (γ) of 10.0. Convergence was assessed by monitoring the time evolution of the collective variables, and was indicated by the absence of systematic drift and large fluctuations over the final portion of the trajectories. The biased simulations were then post-processed using reweighting procedures to reconstruct unbiased ensemble properties and free-energy landscapes. Simulations were conducted using GROMACS in combination with the PLUMED 2.9.3 enhanced sampling plugin.^61, 62^

### 2.3. CG MD Simulations

We performed CG MD simulations using GROMACS 2024.4.^45^ Peptoids were modeled with the CG force field developed in this study, while water was represented using the MARTINI 3 water model.^33^ Initial CG configurations of peptoids were converted from their AA structures using martinize2.^63^ Peptoids were solvated in cubic water boxes ranging from 6 × 6 × 6 nm³ to 12 × 12 × 12 nm³, consistent with the AA system setup. In simulations of multiple chains probing multi-chain aggregation and supramolecular self-assembly, 50 trimer peptoids were randomly placed in a 12.0 × 12.0 × 12.0 nm³ cubic box. The number of water molecules was adjusted to achieve a density of 1.0 g/cm³, ranging from ∼1800 to 14,000 for the single-chain systems in cubic boxes, and ∼12000 for the multiple-chain systems. For systems with net charges, Na⁺ or Cl⁻ ions were added to maintain charge neutrality. Subsequently, forces on beads in excess of 1000 kJ/(mol·nm) were eliminated by the steepest descent energy minimization. Initial atom velocities were assigned from a Maxwell-Boltzmann distribution at 300 K and 1 bar. Simulations began with a 4 ns equilibration under the NPT ensemble, using a velocity rescaling thermostat with a time constant of 1.0 ps and a Berendsen barostat with a time constant of 4.0 ps and a compressibility of 3 × 10⁻⁴ bar⁻¹.^50, 64^ All production runs were carried out in the NPT ensemble at 300 K and 1 bar for 500 ns, employing a velocity rescaling thermostat with a time constant of 1.0 ps and a Parrinello-Rahman barostat with a time constant of 12.0 ps and a compressibility of 3 × 10⁻⁴ bar⁻¹.^65^ For longer chains (e.g., n = 50), the Berendsen barostat was found to provide more stable pressure coupling. A leap-frog integrator was applied with a 10 fs time step.^53^ Non-bonded interactions were treated using the Verlet cutoff scheme with periodic boundary conditions, employing a 1.1 nm cutoff for both electrostatics (εᵣ = 15) and van der Waals forces.^33^ Notably, the *nrexcl* parameter was set to 3 to remove nonbonded interactions between beads separated by up to three bonds, which helps prevent numerical instabilities arising from the overlap of bonded and nonbonded interactions. Importantly, the underlying source of this instability is the substantially stronger bond-angle potentials in peptoids compared to peptides. Trajectory coordinates were saved every 1000 steps in compressed format, while positions, velocities, and forces were not recorded separately. For single-peptoid systems, simulation speeds ranged from 2580.13 to 15990.32 ns/day, representing an estimated 57-fold speedup relative to the corresponding all-atom simulations. To facilitate reproducibility, all input files necessary to perform simulations with our CG model are provided in the *<u>Supporting Information</u>*.

## 3. Model Development

### 3.1. CG Mapping Strategy

Bottom-up parameterization starts from atomistic reference simulations to construct CG models that reproduce the configurational distributions and thermodynamic behavior of their fine-grained counterparts.^66^ A critical design step in this process is the choice of mapping scheme, which dictates how atomistic degrees of freedom are represented by CG beads. In the MARTINI 3 model, a center-of-geometry (COG) mapping is adopted instead of the conventional center-of-mass (COM) approach.^33^ This “size-shape” concept is specifically intended to preserve the molecular volume relative to AA reference structures.^33^ Combined with the proper selection of MARTINI 3 bead sizes, this strategy results in more realistic molecular packing.

#### 3.1.1. Backbone Mapping

Peptoid backbone dihedrals are highly sensitive to N-substitution, even minor changes in backbone grouping can alter conformational preferences and the cis/trans balance.^67^ Consequently, an appropriate CG mapping must yield unimodel, near-symmetric backbone bonded distributions that can be robustly represented by simple harmonic potentials, while preserving backbone conformations and remaining compatible with MARTINI 3 sidechain mapping. Accordingly, taking NVal as an example, we evaluated several candidate backbone mapping schemes to capture peptoid connectivity while maintaining well-behaved bonded distributions suitable for the bottom-up parameterization, as shown in Figure 1. Among these candidates, Mapping 1 satisfies these criteria, whereas Mappings 2-4 exhibit distinct deviations from unimodality or symmetry.

**Figure 1.**
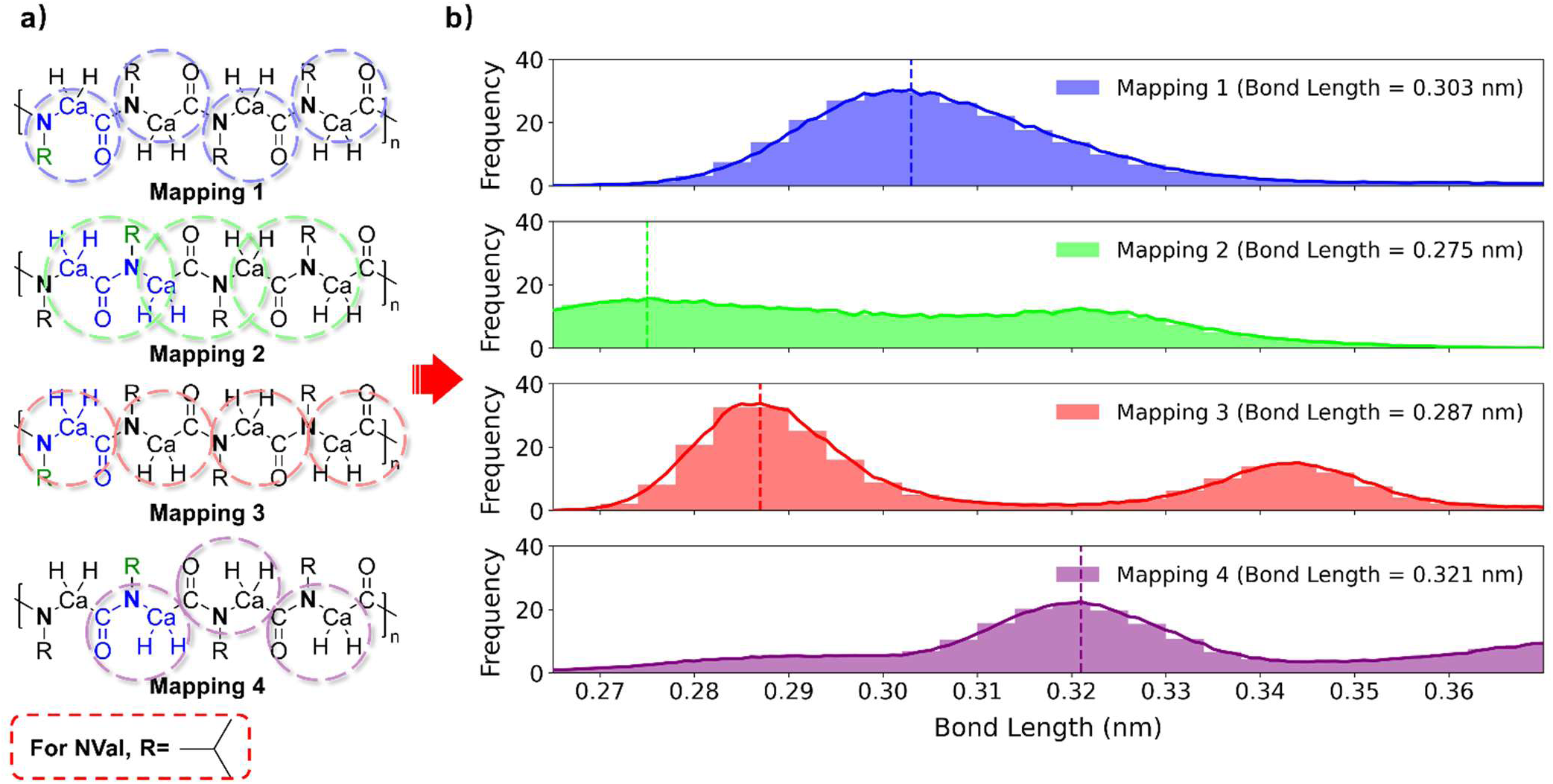
Average backbone bond length distributions of NVal for different mappings. (a) Chemical schematics illustrating the atom selections used to define the geometric centers of CG beads in Mapping Schemes 1-4. Here, R denotes the sidechain, which is represented directly according to the MARTINI 3 definition. Mapping 1’s geometric center is placed along the C–Cα bond, using only the heavy atoms N, Cα, C, and O; Mapping 2’s geometric center is positioned along the C–N bond, using the N, C, and O atoms, half of each CH_2_ group; Mapping 3’s geometric center is placed at the Cα atom, including N, CH_2_ group, C, and O; Mapping 4’s geometric center is shifted closer to the nitrogen atom, using the N, C, and O atoms along with CH_2_ group bonded to it. (b) Bond length distributions of NVal for the four mapping schemes. The *x*-axis represents bond length (nm), while the *y*-axis indicates frequency. Bond length distributions were obtained from four homopeptoid sequences and averaged to produce the shown profiles.

Mapping 2, inspired by previous work, defines each CG backbone bead across two residues as a PA bead centered at the midpoint of the C–N backbone bond.^29^ Gao and Tartakovsky observed that introducing such a nonstandard MARTINI bead yielded a simpler distribution function for the PA–PA interaction.^25^ The PA bead encompasses the backbone N, Cα, and O atoms along with half of each backbone CH₂ group. However, in our COG case, the mapping produced a double-peaked bond length distribution with an equilibrium distance of ∼0.275 nm, suggesting artificial conformational frustration. Relocating the mapping center toward the backbone nitrogen (Mapping 4) did not resolve this issue, as the bond length distribution remained bimodal. Since the attachment site of the backbone hydrogen differs between peptides and peptoids, we tested Mapping 3 for shape fidelity by locating the geometric center at Cα and defining it by N, Cα, Hα, C, and O. While this scheme better preserved molecular volume, the bond length histogram showed multimodality, likely reflecting the contribution of hydrogen atoms, which could lead to asymmetric positional fluctuations and shifting in the geometric center. To address this, we refined Mapping 1 by excluding the hydrogens and retaining only the heavy atom scaffold of each residue. This refinement yielded a single, well-defined bond length distribution and a corresponding CG potential. A key advantage of this refinement is that the bonded terms are fully compatible with MARTINI and avoid the need for de novo fitting from all-atom reference simulations. For bead type assignment in MARTINI 2, the backbone particle class was secondary-structure dependent and explicitly coupled with the BB-based bonded parameters.^68^ In contrast, we adopt the current framework, which employs a uniform representation wherein all BB beads are consistently modeled as P2, independent of the underlying secondary structure.^32^

#### 3.1.2. Sidechain Mapping

Sidechain mappings were constructed to follow the standard bead definitions, sizes, and mass assignments provided by the MARTINI 3 forcefield, enabling direct reuse of the existing chemical ontology and ensuring full compatibility with other MARTINI 3 biomolecular components. Each peptoid residue was mapped using a COG scheme consistent with MARTINI 3 guidelines, with hydrophobic, polar, aromatic, and charged residues represented by their corresponding bead classes (see Figure 2 for full residue mapping schemes). Because MARTINI 3 already provides chemically specific bead labels for a broad set of functional groups, no new bead types were introduced in this work.

**Figure 2.**
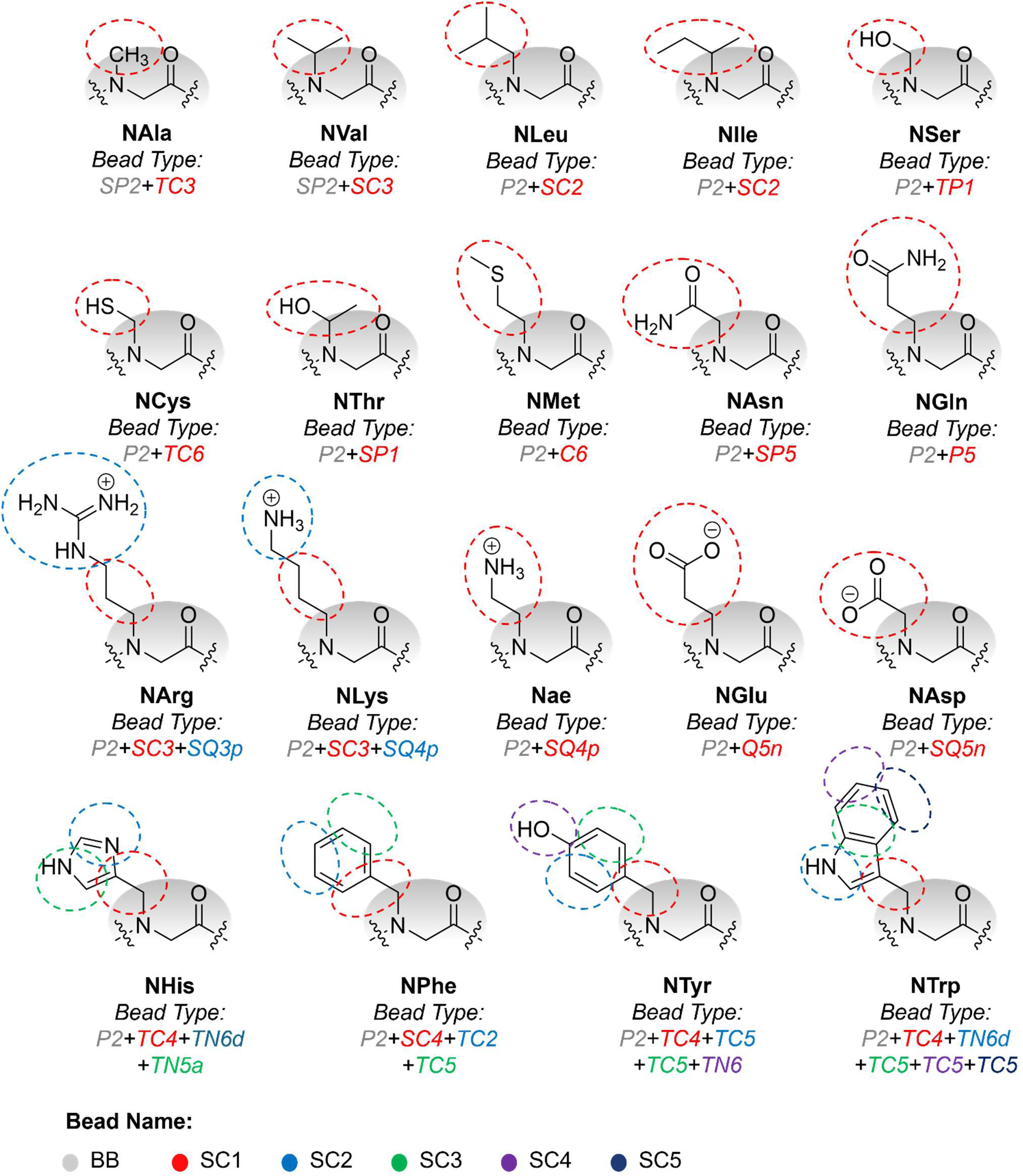
CG representation of the 19 canonical peptoid side chains developed in this work. For each residue type, the corresponding MARTINI 3 bead assignments are shown, with the backbone mapped to the P2 bead and the side-chain moieties mapped to chemically appropriate polar, apolar, charged, or aromatic bead classes.

For most residues, the MARTINI 3 sidechain definitions yielded well-behaved bonded distributions when coupled to the unified peptoid backbone. However, in a small subset of residues, the inclusion of explicit hydrogen atoms in the SC1 bead caused asymmetric COG shifts that resulted in multimodel BB–SC1 bond distribution and poor harmonic fitting. For these residues, refinements were introduced to the hydrogen representation within SC1 beads to enforce well-defined, unimodal BB–SC1 bond distributions consistent with harmonic potential behavior (*<u>See Supporting Information for details</u>*). This design preserves the modularity and transferability of peptoid sidechains across homo- and heteropeptoid sequences while ensuring seamless integration with the MARTINI 3 nonbonded interaction matrix.

### 3.2. Bonded Interaction Parameterization

#### 3.2.1. Bond, Angle, and Dihedral Parameterization

The bonded parameters were obtained via direct Boltzmann inversion, which derives the mean force for a given degree of freedom.^69, 70^ In the present work, initial estimates of the CG bonded interaction potentials were generated by DBI of distribution functions extracted from PBMetaD-enhanced all-atom reference simulations of 4mer homopeptoids, based on the defined atom-to-bead mapping. To ensure the compatibility and transferability of the developed forcefield, we parameterized only interactions involving the backbone and the first side-chain bead bonded to it, while all other interactions between side chains were directly adopted from MARTINI 3 without modification (Figure 3). To be specific, we parametrized BB–BB bonds, BB–SC1 bonds, BB^-^–BB–BB^+^ angles, BB–BB^+^–SC1^+^ angles, SC1–BB–BB^+^ angles, SCX–SC1–BB angles (only if this angle exists), BB^-^–BB–BB^+^–BB^++^ dihedrals, BB^-^–BB–BB^+^–SC1^+^ dihedrals, SC1^-^–BB^-^–BB–BB^+^ dihedrals, and SC1–BB–BB^+^–SC1^+^ dihedrals for peptoids.

**Figure 3.**
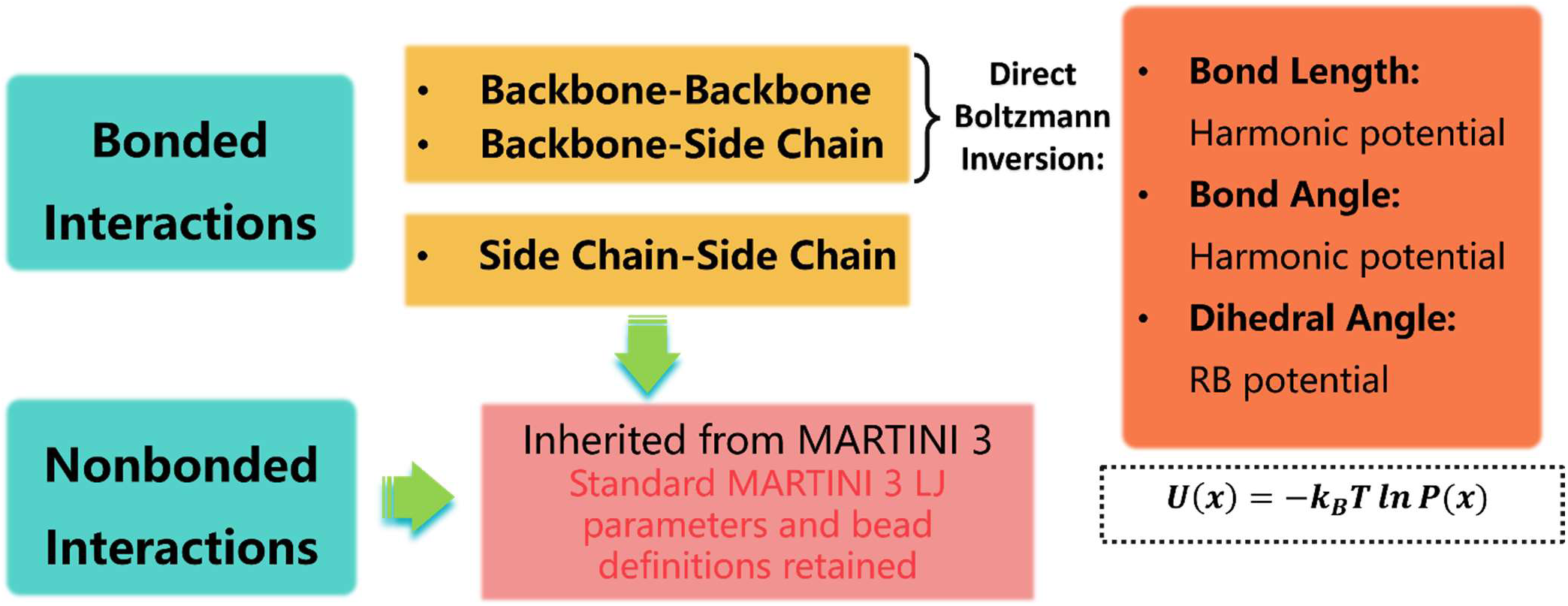
**Schematic overview of bottom-up parameterization in the CG peptoid model.**

The bonded potential is given by:

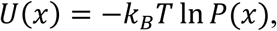

where *P*(*x*) denoting the probability distribution of variable 𝑥. In practice, this involves inverting the probability distribution of the variable of interest, which may correspond to a bond length, bond angle, or torsion angle. To be specific, we use a harmonic potential (Func. Type 1 in GROMACS) for bonds, defined by the equilibrium bond length *bij* (nm) and the force constant *kij* (kJ·mol⁻¹·nm⁻²). Bond stretching between two covalently bonded beads *i* and *j* is thus represented by a harmonic potential:

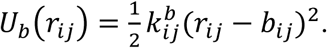

The bond–angle vibration among a triplet of beads *i–j–k* is also represented by a harmonic potential (Func. Type 1 in GROMACS), having the same form as bond stretching:

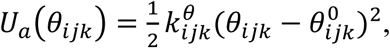

where *θijk* is the bond-angle value, *k^θ^ijk* is the corresponding force constant. Unlike bond length, the equilibrium angle *θ^0^ijk* is determined by harmonic fitting rather than by the most probable value.

Dihedral torsions involving four beads *i–j–k–l* are modeled using the Ryckaert–Bellemans potential (Func. Type 3 in GROMACS), expressed as:

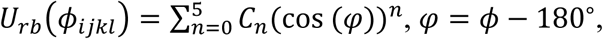

where *Cn* are empirical coefficients.^71^

Due to the enhanced backbone flexibility of peptoids, several DBI-derived distributions exhibited broad or noisy features. The raw DBI profiles were fitted directly to the corresponding functional forms described before. This fitting procedure smooths statistical noise and ensures numerically stable bonded potentials for subsequent CG simulations.

#### 3.2.2. Unified Backbone Bonded Potentials

As peptoids’ backbones generally lack stable secondary structures, their bonded parameters are expected to arise from local chemical environments rather than collective conformational restraints.^72^ Despite sidechain-specific influences, several backbone bonded distributions showed only weak variation across the 19 residues. To ensure compatibility with MARTINI 3, which adopts a unified backbone representation, and to enable transferability across heterogeneous sequences, we derived a unified strategy for those peptoid backbone potentials that is dominated by the shared backbone skeleton rather than the chemical identity of the sidechain. For example, all backbone bond stretching terms (BB–BB) were derived via simple averaging across residues, yielding an equilibrium bond length of 0.310 nm and a force constant of 11207.15 kJ·mol⁻¹·nm⁻² with a standard deviation negligible in magnitude. The same procedure was applied to bond angles (i.e., BB^-^–BB–BB^+^ and SC1–BB–BB^+^ angles) and dihedrals (i.e., BB^-^–BB–BB^+^–BB^++^, BB^-^–BB–BB^+^–SC1^+^, SC1^-^–BB^-^–BB–BB^+^, and SC1–BB–BB^+^–SC1^+^ dihedrals).

Since the dihedral fitting function is represented by coefficients rather than explicit bond angles, point-wise fitting was performed, and the averaged coefficients were taken as the force field parameters for these four different types. All systems here are homopolymer chains, and residue-specific variations were not considered. The detailed residue-specific parameters are summarized in Tables S1-S5 and Figures S1-S4 in the Supporting Information.

A special case is the BB–BB^+^–SC1^+^ angle, which showed substantially stronger and more residue-dependent behavior than other backbone angles, consistent with the geometric constraints imposed by peptoid cis/trans isomerization (Figure 4). For this angle, the DBI-derived force constants were roughly an order of magnitude larger than those of other angles, with equilibrium values showing significant residue-to-residue variation. Because simple averaging produced overly broad or poorly defined parameters, we applied *k-means* clustering, a vector quantization method, to classify the angle distribution because it exhibited distinct trends.^73^ This algorithm partitions *n* observations into *k* clusters, assigning each observation to the cluster defined by the nearest centroid. Accordingly, the 18 residues were grouped into five clusters, and for each category, both the equilibrium angle and the fitted force constant were averaged. The Nae residue was parameterized independently for subsequent validation. Together, the unified backbone bonded interactions, combined with the clustered BB–BB^+^–SC1^+^ angle definitions, yield a transferable and physically grounded bonded interaction framework suitable for both homo- and heteropeptoids with full MARTINI 3 compatibility. Notably, the BB–BB^+^–SC1^+^ angle is the only across-residue bonded term that requires residue-specific parameterization, whereas other residue-dependent bonded terms are limited to within-residue interactions. This design enables straightforward extensibility: new residues can be incorporated by mapping their atomistic angle distributions to the nearest existing cluster, or by defining a new cluster when distinct behavior emerges.

**Figure 4.**
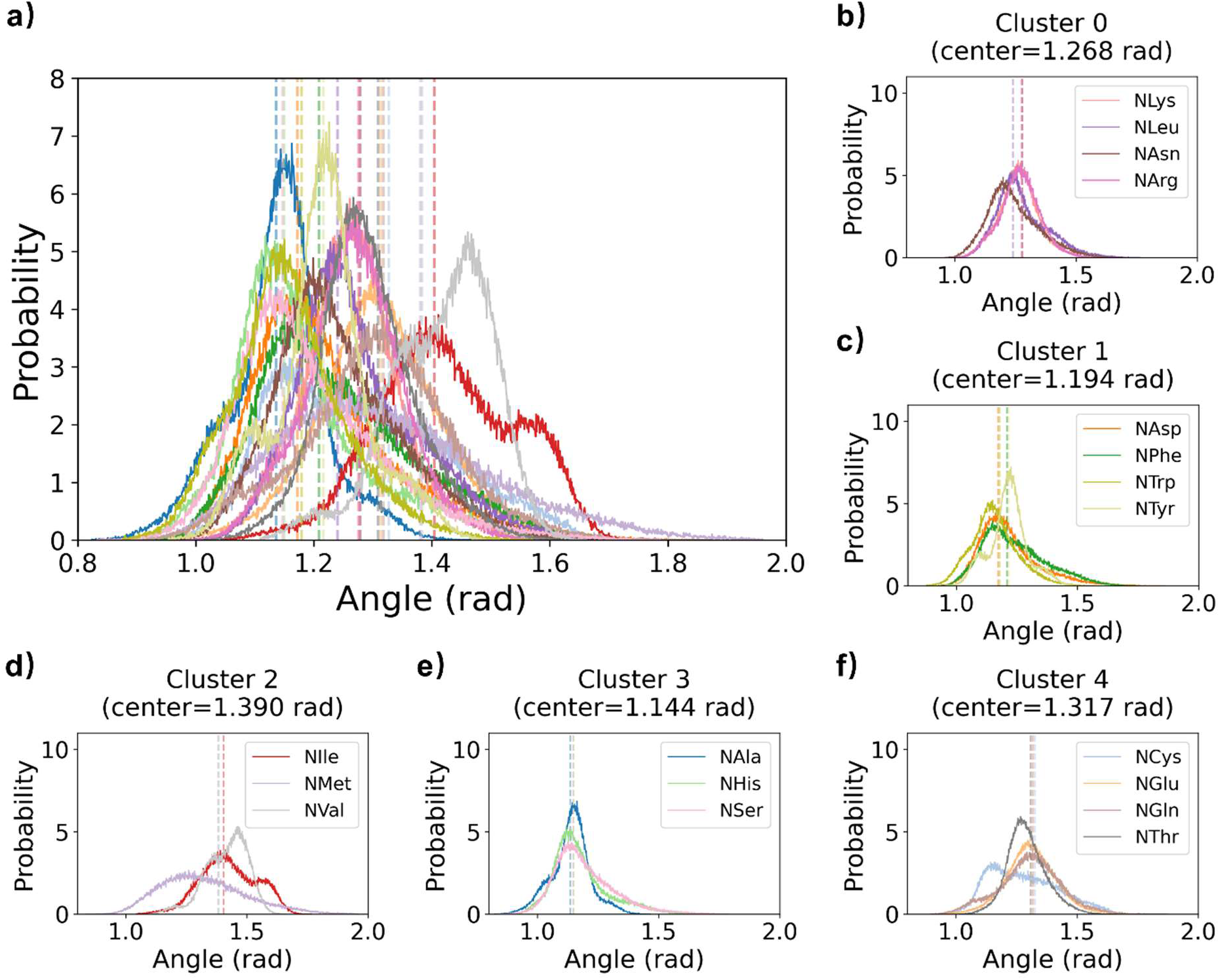
Residue-specific BB–BB^+^–SC1^+^ angle distribution in the database mapped from AA trajectory. Based on residue specific angle distribution, 18 residues were split into 5 for BB–BB^+^–SC1^+^ angle definition. (a) Overall BB–BB^+^–SC1^+^ angle distribution for all residues in the database. (b–f) BB–BB^+^–SC1^+^ angle distributions for the five residue clusters.

The resulting backbone bonded parameters are summarized in Table 1.

**Table 1.** CG Backbone Bonded Parameters.

| Bond Type |  | Bond Length (nm) | Force Constant (kJ/mol·nm<sup>2</sup>) |
| --- | --- | --- | --- |
| BB–BB<sup>+</sup> |  | 0.31 | 11207.150 |
| Bond Angle Type |  | Bond Angle (rad) | Force Constant (kJ/mol·rad<sup>2</sup>) |
| BB<sup>−</sup>–BB–BB<sup>+</sup> |  | 2.31 | 23.480 |
| BB<sup>−</sup>–BB–SC1 | Cluster 0 | 1.27 | 330.215 |
|  | Cluster 1 | 1.19 | 339.842 |
|  | Cluster 2 | 1.39 | 224.343 |
|  | Cluster 3 | 1.14 | 327.513 |
|  | Cluster 4 | 1.32 | 189.608 |
| SC1–BB–BB<sup>+</sup> |  | 2.46 | 18.870 |

### 3.3. Nonbonded interactions

MARTINI 3 introduces more than 800 bead types together with a thoroughly recalibrated interaction matrix, and the number of possible interaction levels has been extended from 10 to 22.^74^ Our mapping scheme developed here uses only standard MARTINI 3 bead types, thereby enabling direct use of the standard non-bonded framework without further modification. This design ensures fully compatible with the broader MARTINI 3 ecosystem and maximizes transferability across biomolecular and materials simulations.

## 4. Validation of the CG Models

To assess the accuracy and transferability of the CG peptoid model, we performed a series of validation tests spanning multiple structure and thermodynamic observables. These benchmarks include polymer scaling behavior, backbone dihedral free-energy landscapes, dimerization potentials of mean force, and multi-chain aggregation of heterotrimers. Each test directly compares CG predictions with corresponding atomistic reference simulations, enabling us to evaluate how well the model reproduces key conformational and assembly features across length and time scales relevant to peptoid behavior.

### 4.1. Polymer Scaling Behavior (Rg)

We first evaluate the structural behavior of isolated peptoid chains in water by comparing the Rg from CG and AA simulations for eight representative residues over 500 ns simulations. All chains were initialized in the all-trans configuration. Rg characterizes the spatial distribution of a polymer’s mass around its center of geometry, providing a global measure of chain compactness.^75^ As a key descriptor of polymer size, Rg provides a direct benchmark for assessing whether the CG model preserves the global dimensions of the all-atom system. Figure 5a presents the Rg value of eight representative residue chains (NAla, NCys, NAsp, NLys, NThr, NVal, NPhe, NTrp) with polymerization degrees N = 4, 6, 8, 10, 15, and 20, obtained from AA and CG models under identical conditions (T = 300 K and P = 1 bar). For short chains (N ≤ 10), the CG model reproduces the AA Rg values with good fidelity (below 30%) across all residue types except NAsp.^76^ As chain length increases, however, the deviations become more pronounced, particularly N = 20. Part of this discrepancy may reflect limitations of AA simulations for longer chains, where convergence becomes more challenging and the equilibrium cis/trans distribution may not be fully sampled. In addition, as the four-residue parameterization emphasized terminal environments, while longer chains involve interior residues with different flexibility and local environments, amplifying the deviation. Taken together, the CG model reproduces the expected scaling trends and global size of peptoid chains, with particularly strong agreement in the short-chain regime and physically interpretable deviations at larger chain lengths.

**Figure 5.**
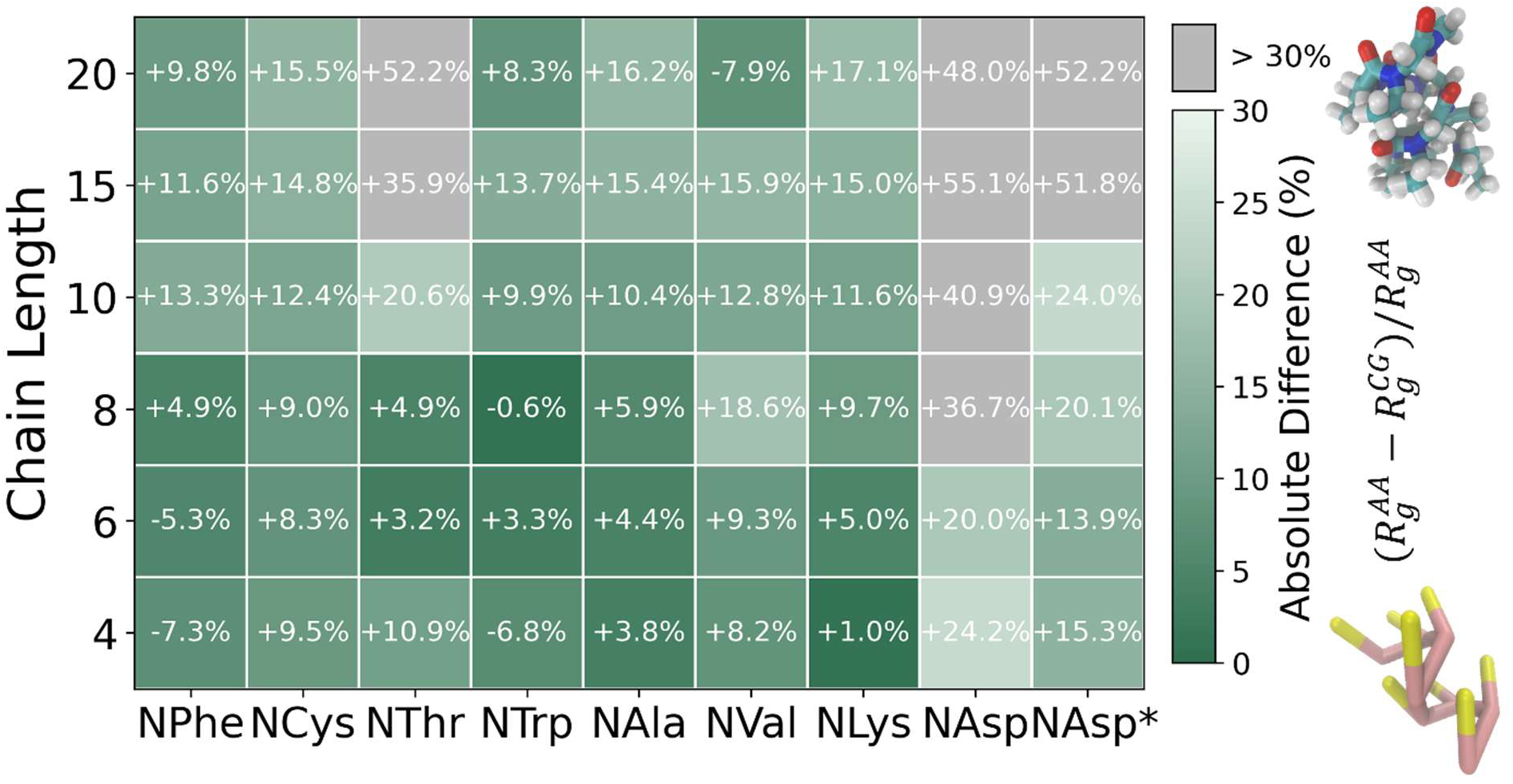
Radius of gyration of homochains from AA and CG models at different chain length. **N.** Eight representa-tive residues were selected to cover major side-chain types: NAla (reference), NVal (branched), NCysd (sulfur), NAsp (acidic), NLys (basic), NThr (polar), NPhe (aromatic), and NTrp (aromatic heterocycle). NAsp* denotes the residue with refined improper dihedrals. The percentage values represent the relative difference between the two models, calculated as (𝑅*_g_*^AA^ − 𝑅*_g_*^CG^)/𝑅*_g_*^AA^. Green denotes differences with [- 30%, 30%] and grey denotes differences > 30% or < - 30%.

A notable exception is NAsp, for which the CG model consistently predicts markedly smaller Rg values than the corresponding AA simulations even at short chain lengths. This behavior is likely related to differences in charge representation between the AA and CG models. In the AA model, the −1 charge of NAsp is distributed over multiple atoms, whereas in the CG model it is assigned to a single bead, which can enhance electrostatic interactions and promote more compact conformations. This also happens to NLys group. However, in the all-atom model, charges on NH3^+^ group are more localized. So, the differences between CG and AA are smaller.

These observations are consistent with previous reports that simulations with MARTINI 3 on average underestimate Rg by nearly 30% without tailed bonded parameters for a given class of disordered chains and suggest a rescaling factor for increased protein-water interactions.^76^ In response, the MARTINI developers concluded that the absence of optimized bonded terms was the primary cause of conformational collapse and therefore refined the bonded parameterization for intrinsically disordered proteins (IDPs), giving rise to the MARTINI3-IDP model.^77^ A key modification was the introduction of BB^-^–BB–SC–BB^+^ improper dihedrals, which reflect the intrinsic sp³ tetrahedral geometry of the Cα atom and improve the agreement with atomistic conformational ensembles. To assess whether a similar correction is required for peptoids, we compared the improper dihedral distributions obtained from our AA reference simulations with those generated by the CG model for the representative residues (Figure S5). For nearly all residues examined, the CG improper dihedral distributions closely matched their AA counterparts, indicating that the existing bonded terms already capture the relevant backbone geometry without requiring additional improper interactions. NAsp was the only exception, exhibiting a clear deviation between AA and CG improper dihedrals, consistent with its anomalously compact Rg. To address this issue, residue-specific improper dihedrals were introduced. For NAsp*, this refinement led to a noticeable improvement in short chains. However, for longer chains, the improvement is limited, as intrinsic intramolecular interactions dominate, resulting in increased deviations from the defined improper geometry and larger discrepancies in Rg. Similar improper terms were also introduced for NGlu and Nae, where charged beads are directly connected to the backbone.

### 4.2. Backbone Dihedral Free Energy Surface

To evaluate whether the CG model captures backbone conformational preferences, we examined the free energy surface (FES) of adjacent backbone dihedral angles (ψ1, ψ2) for a subset of 6-residue homopeptoids. For each residue type, we compute (i) unbiased AA dihedral distributions, (ii) corresponding PBMetaD-enhanced AA distributions, and (ii) CG distributions (Figure 6).

**Figure 6.**
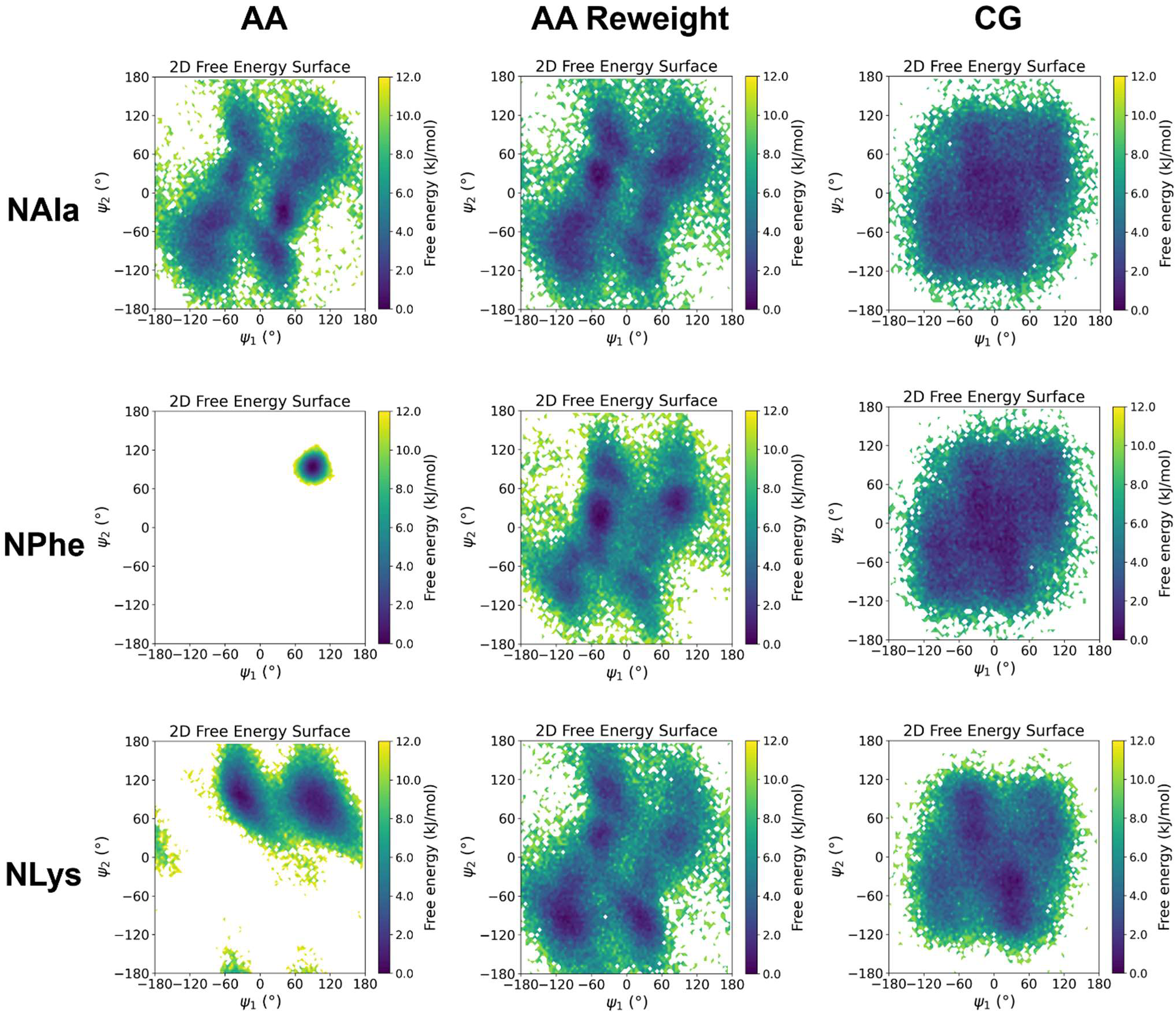
Free energy surfaces of backbone dihedrals for three representative 6-residue homochains, obtained from AA MD, reweighted PBMetaD AA MD, and CG simulations. ψ_1_, ψ_2_ denote two adjacent backbone dihedral angles.

Unbiased AA simulations exhibit narrow and often incomplete FES, reflecting insufficient sampling of slow cis/trans isomerization and associated backbone rearrangements. In contrast, the PBMetaD-enhanced AA simulations achieve substantially broader and more continuous coverage of the dihedral landscape, providing a converged reference for backbone conformational sampling. The CG free energy surfaces display comparable coverage to the PBMetaD-enhanced AA results and reproduce the overall topology of the sampled conformational space across all examined residue types, ranging from the simplest side chains to charged and aromatic residues.

In particular, for the ring-containing residue (i.e., NPhe), where unbiased AA sampling is especially restricted, the CG model reproduces the broad backbone conformational regions identified in the reweighted PBMetaD AA reference. As expected from the coarse-graining procedure, multiple atomistic microstates are projected onto a reduced set of CG degrees of freedom. As a result, the CG model, which employs a uniform backbone parameter set with position-independent potentials, produces broadly similar free-energy surfaces across different residues. While the overall topology of the conformational space is well reproduced, finer residue-specific angular coupling present at the atomistic level is partially averaged out due to the coarse-grained representation. Overall, the CG model accurately captures the ensemble-averaged backbone conformational landscape defined by the converged AA free energy surfaces, while highlighting that finer residue-independent angular correlations would require more detailed or higher-order backbone representations in future CG forcefield developments.

### 4.3. Dimerization Potential of Mean Force

We evaluated the tendency of peptoid chains to associate in water by computing the potential of mean force for dimerization of four representative homotetramers: NVal (hydrophobic), NCys (polar), NArg (cationic), and NPhe (aromatic). Umbrella sampling simulations were performed at T = 300 K and P = 1 bar using the COM separation between the center backbone beads of the two chains as the reaction coordinate, as defined by the all-atom to CG mapping scheme. Sampling was performed over the range *d* = 0.5-3.5 nm with increments of 0.1 nm. In each window, harmonic biasing potentials with force constants of 1000-2000 kJ·mol⁻¹·nm⁻² were applied, followed by 10 ns of production simulation. The unbiased PMFs were reconstructed from the biased trajectories using the weighted histogram analysis method (WHAM) implemented in GROMACS.^78, 79^

The PMF curves reported in Figure 7 show that the CG model reproduces the key features of the corresponding AA reference profiles, including the locations of free-energy minima and the overall barrier shapes. For each residue, the CG dimerization PMF is in good agreement with the AA reference, indicating that the model captures residue-specific association behavior.

**Figure 7.**
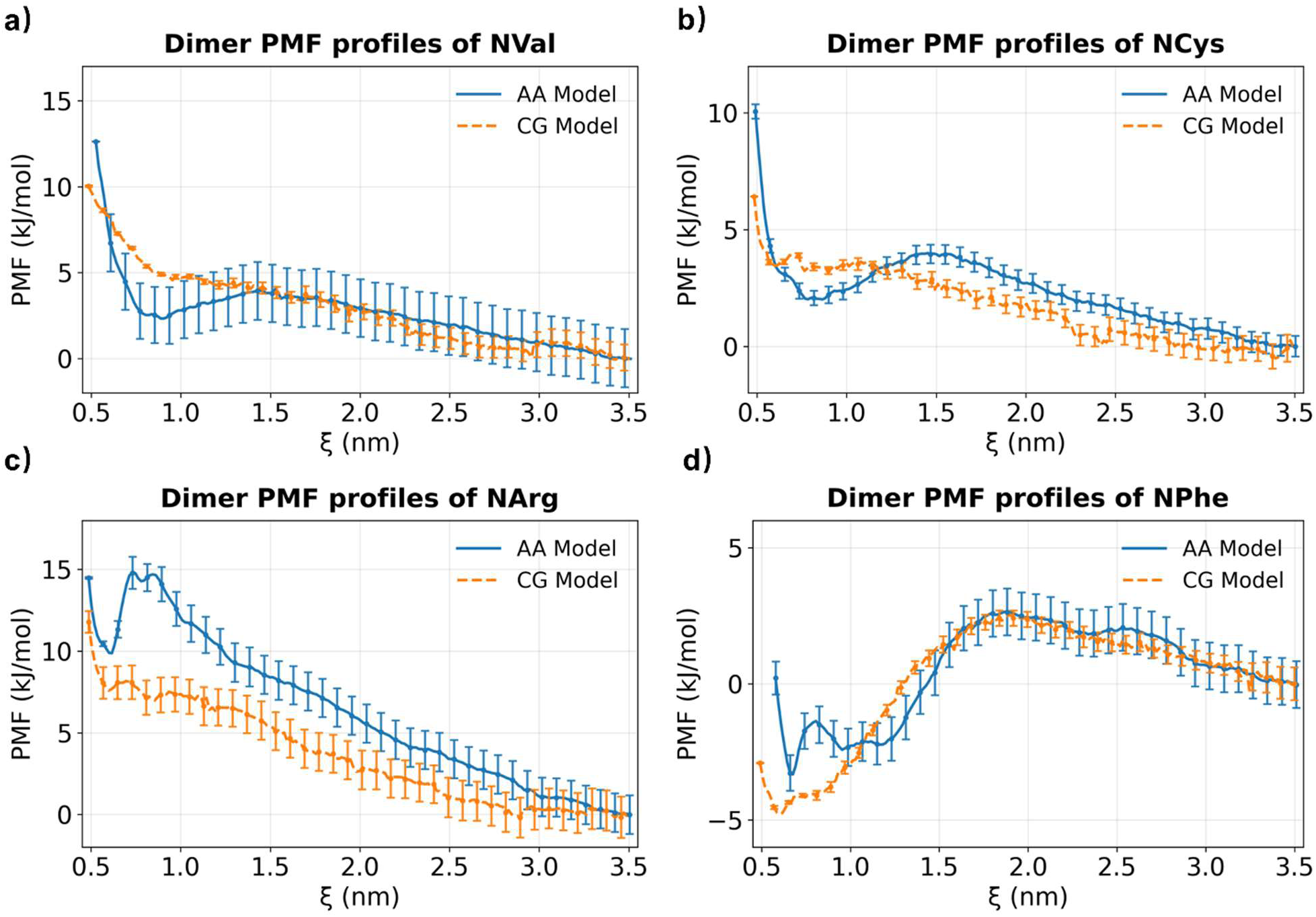
Dimerization PMSs for four homo-tetramers (panels a-d). PMF curves are computed using umbrella sampling and WHAM, and uncertainties are estimated by 200 rounds of bootstrap resampling. The free energy at the largest chain separations was set to zero, serving as the reference state corresponding to two non-interacting chains.^80^

Quantitatively, the differences between AA and CG PMFs are small. Defining the dimerization free energy for NPhe, ΔFdimer, as the difference between this reference value and the minimum of the free-energy profile located at *d* ≈ 1.0 nm, we find ΔFdimer^CG^=−(2.932 ±0.188) kJ·mol⁻¹ and ΔFdimer^AA^=−(2.280 ± 0.687) kJ·mol⁻¹, which differ by only 0.65 kJ·mol⁻¹. Applying the same definition to the remaining three systems yields unfavorable dimerization free energies that nevertheless remain agree within 1.69 *kB*T at T=300 K. Across the full separation range, deviations remain within the expected accuracy of MARTINI-level coarse-graining.^33^ Minor discrepancy appears at short separations; for example, the shallow local maximum at d ≈ 0.7 nm in the AA NArg system arises from specific atomistic interactions, including hydrogen bonding, which are not explicitly represented in the CG model.Overall, these results demonstrate that the CG model reliably reproduces AA dimerization thermodynamics on a residue-by-residue basis.

### 4.4. Sequence-Dependent Early-Stage Aggregation Behavior

Peptide nanowires hold promise for applications in biomimetic electronics, nanoparticle joining, and cell culture scaffolds.^81–83^ Peptoids further extend this potential, as their unique structural features and functional groups broaden molecular versatility and may enable novel applications. Motivated by these considerations, our final test assessed whether the CG model could capture peptoid self-assembly behavior. Although most experimental studies have focused on long-chain peptoids, the ordered supramolecular assemblies they form are challenging to simulate due to the long time scales required.^84–86^ In contrast, Castelletto and co-workers investigated the assembly in water of short heterotrimers containing phenyl (NPhe) and lysine analogs (NLys/Nae), specifically NPhe–Nae–NPhe (FXF), NPhe–NLys–NPhe (FKF), Nae–NPhe–NPhe (XFF), and NLys–NPhe–NPhe (KFF).^87^ Cryo-TEM revealed that only FXF assembled into linear nanowires, whereas the other three formed globular structures. Notably, the previously developed MoSiC-CGenFF-NTOID model successfully reproduced these experimental findings atomistically by identifying FXF as an outlier relative to FKF, KFF, and XFF.^22^ This result supports the model’s capability in predicting aggregation trends in multi-chain peptoid systems and lays the groundwork.

We simulated 50-chain systems of FKF, FXF, KFF, and XFF heterotrimers in water for 500 ns. Nae (X) was parameterized separately for bonds and angles and included in the CG model. All sequences displayed self-assembly behavior, leading to the formation of long-lived clusters. In the analysis, each molecule was represented as a node, with edges defined by intermolecular heavy-atom contacts within 5.0 Å; clusters were identified as connected subgraphs containing three or more nodes. To account for the non-equilibrium and history-dependent nature of clustering, three independent trajectories were performed, and only the final 200 ns of each trajectory were analyzed. In Figure 8a, we present molecular visualizations of representative clusters for four heterotrimers. All sequences form clusters ranging from 3 to 50 molecules, with continuous association and dissociation events observed throughout the simulation. Consistent with trends observed in AA simulations,^22^ FXF exhibited the smallest mean cluster size (43.187 ± 0.815) compared to FKF (45.851 ± 2.229), KFF (46.357 ± 0.220), and XFF (43.464 ± 0.259), with uncertainties representing standard errors across the three runs (Figure 8b). While the relative ordering agrees with AA results, the CG model systematically predicts larger cluster sizes, reflecting the known tendency of the CG models to enhance aggregation. Nevertheless, the CG model captures the correct sequence-dependent aggregation trends across systems.

**Figure 8.**
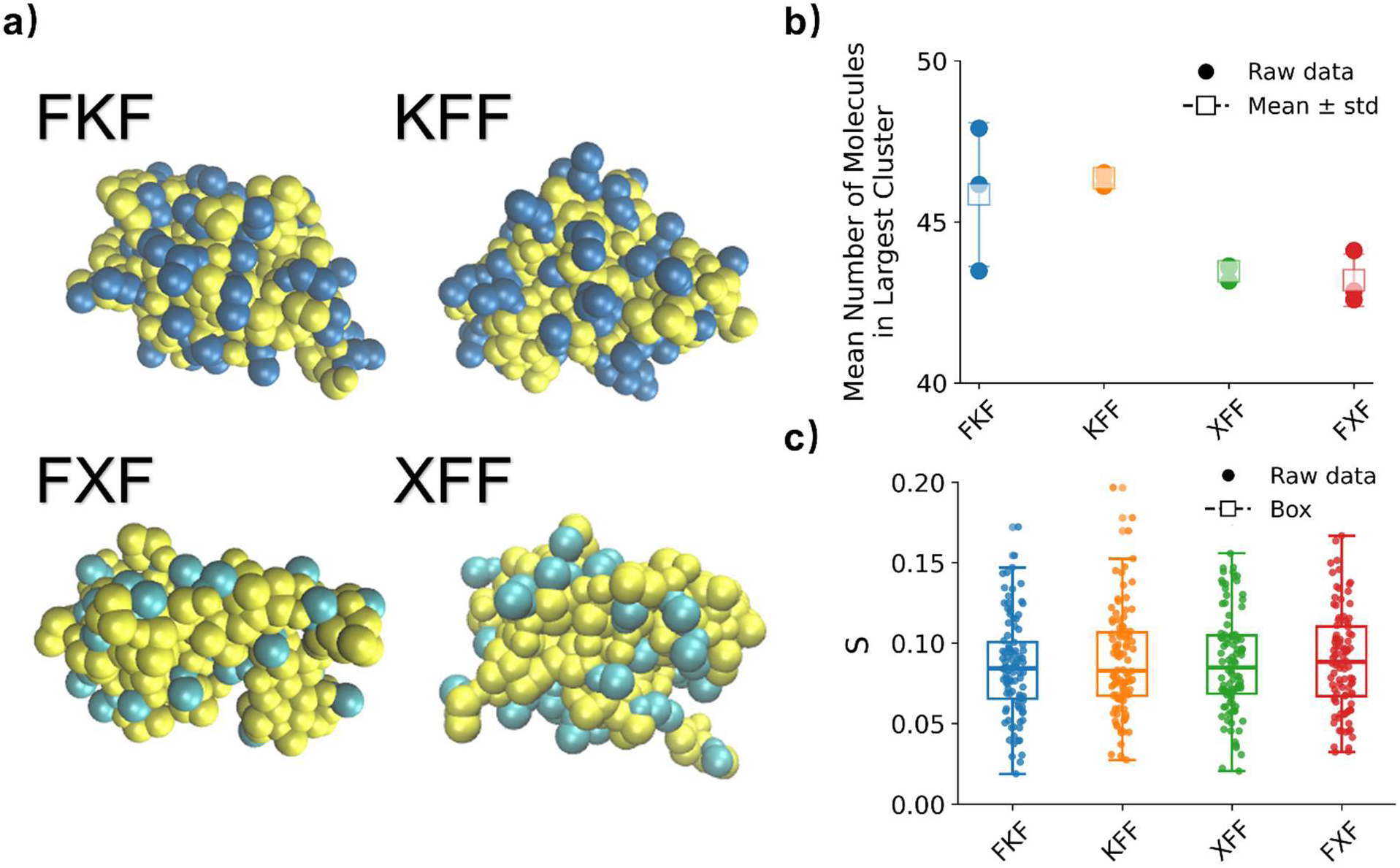
Self-assembly and orientational ordering of heterotrimers system. (a) Visualization of four heterotrimers observed in simulations of 50 peptoid chains in water, colored by residue type. (b) Largest-cluster size, results from three independent simulations are shown as dots, and averages with error bars are shown as squares. (c) Orientational order parameter (S) of the four sequences. Each data point represents the S value from an individual simulation frame, and the box plots summarize the overall distributions across trajectories.

To investigate the sequence packing patterns within clusters, we also examine the orientational order parameters (*S*),which quantifies peptoids alignment in the bulk.^88^ At each time frame *t*, the instantaneous alignment tensor was defined as:

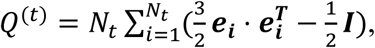

where *Nt* is the number of molecules at time *t*, 𝑒_i_ denotes the unit orientation vector of aromatic ring *i*, and *I* is the identity matrix. The instantaneous scalar order parameter was then defined as the largest eigenvalue of *Q*(t), denoted *St*. The trajectory-averaged order parameter was then obtained by averaging over all sampled frames, approaching 1 for perfectly aligned ring orientations and 0 for random orientations.^89^

As shown in Figure 8c, all sequences exhibit relatively low orientational order, consistent with the early-stage aggregation regime sampled in these simulations. Although FXF shows a slightly higher mean S, the distributions across sequences substantially overlap, and no statistically significant difference are observed among sequences (Kruskal-Wallis test, *p* = 0.129, *n* = 3 per sequence). This suggests that global orientational alignment is comparable across sequences under the conditions studied. Together, these results indicate that while the CG models capture general aggregation behavior, pronounced or sequence-specific ring alignment does not emerge at this stage of assembly.

## 5. Conclusion

In this work, we developed a bottom-up CG peptoid forcefield encompassing 19 residues using a unified and chemically consistent mapping scheme fully compatible with the MARTINI 3 framework. Extensive enhanced sampling at the AA level was employed to generate converged reference data for slow backbone ω-dihedral cis/trans isomerization, enabling systematic parametrization of bonded interactions via direct Boltzmann inversion, while nonbonded interactions were largely adopted from the standard MARTINI 3 parametrization with targeted refinements to ensure peptide-to-peptoid transferability.

The resulting forcefield was rigorously validated against a diverse set of atomistic benchmarks, including polymer scaling behavior, backbone dihedral free-energy surfaces, dimerization potentials of mean force, and early-stage heterotrimer aggregation. Across these tests, the CG model shows consistently good agreement with AA reference data for both structural and thermodynamic observables, demonstrating that a unified MARTINI-compatible backbone description is sufficient to capture the key features of peptoid backbone flexibility, sequence-dependent conformational behavior, and early-stage assembly. Importantly, this framework retains the computational efficiency of CG simulations while enabling chemically specific and transferable modeling of sequence-defined peptoid systems.

The present model, implemented with martinize2 workflow (https://github.com/MZhaoLab/polypeptoid_cg_model), provides a practical and extensible platform for mesoscale simulations of peptoid structures, dynamics, and self-assembly. It also establishes a foundation for future developments of MARTINI 3 models targeting synthetic foldamers and sequence-defined polymers.

## Acknowledgement

This work was supported by the start-up fund from the College of Engineering at The University of Alabama. Computational resources were provided by the University of Alabama High-Performance Computing (UAHPC) facility, supported by the Office of Research & Economic Development and the Office of Information Technology. We also thank the Alabama Supercomputer Authority (ASA) for access to its ASA-X high-performance computing infrastructure, which provides HPC services at no cost to Alabama’s public universities, faculty, staff, and students.

## TOC figure

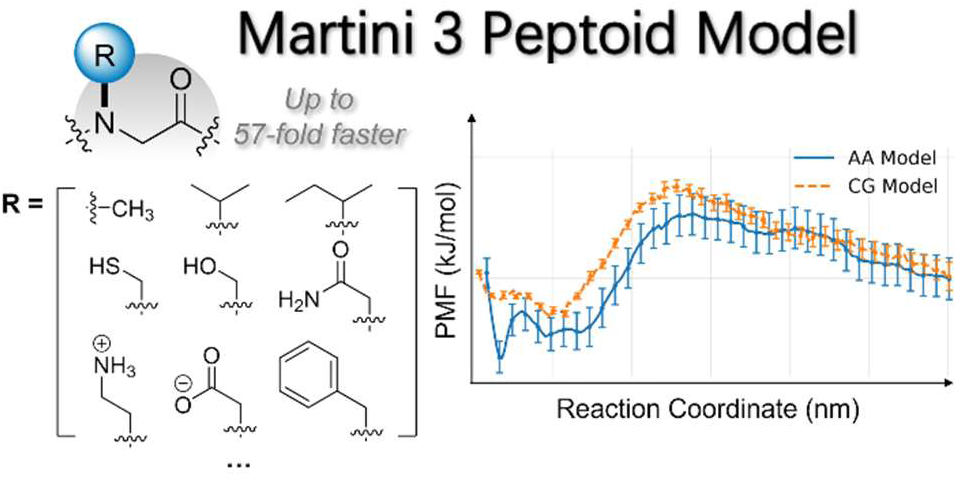

