## Supporting Information for "Extending the MARTINI 3 Coarse-Grained Forcefield to Polypeptoids"

### 1. All-Atom Parameterization of Individual Peptoid Residues

#### 1.1 Bond and Angle Parameters for Each Residue

Before applying the unified mapping scheme, each peptoid residue was individually parameterized at the all-atom level. For completeness, the residue-specific bonded parameters, including the fitted backbone bond lengths, bond angles, and dihedral terms derived from Boltzmann-inversion, are summarized below.

**Table S1.** Bonded Parameters for BB–BB$^+$ Bonds Across 18 Residues.

| Residue Name | BB–BB$^+$ Bong Length (nm) | Force Constant (kJ/mol·nm$^2$) |
|---|---|---|
| NAla (A) | 0.314 | 10607.92 |
| NCys (C) | 0.313 | 9789.05 |
| NAsp (D) | 0.315 | 9074.87 |
| NGlu (E) | 0.311 | 10783.75 |
| NIle (I) | 0.298 | 15904.89 |
| NLeu (L) | 0.308 | 11522.77 |
| NMet (M) | 0.312 | 10928.61 |
| NAsn (N) | 0.313 | 9722.68 |
| NGln (Q) | 0.310 | 11864.65 |
| NSer (S) | 0.317 | 9123.84 |
| NThr (T) | 0.307 | 11412.26 |
| NVal (V) | 0.303 | 13964.13 |
| NLys (K) | 0.312 | 12336.31 |
| NArg (R) | 0.308 | 12574.12 |
| NHis (H) | 0.311 | 10925.28 |
| NPhe (F) | 0.310 | 10370.26 |
| NTyr (Y) | 0.311 | 10972.11 |
| NTrp (W) | 0.312 | 9851.28 |

**Table S2.** Bonded Parameters for BB–SC1 Bonds Across 18 Residues.

| Residue Name | BB–SC1 Bong Length (nm) | Force Constant (kJ/mol·nm$^2$) |
|---|---|---|
| NAla (A) | 0.255 | 14792.67 |
| NCys (C) | 0.352 | 2290.68 |

| | | |
|---|---|---|
| NAsp (D) | 0.364 | 11627.93 |
| NGlu (E) | 0.372 | 4254.34 |
| NIle (I) | 0.311 | 8850.57 |
| NLeu (L) | 0.376 | 13036.93 |
| NMet (M) | 0.395 | 1537.57 |
| NAsn (N) | 0.323 | 4087.19 |
| NGln (Q) | 0.358 | 3453.34 |
| NSer (S) | 0.319 | 5378.40 |
| NThr (T) | 0.299 | 10228.44 |
| NVal (V) | 0.283 | 25425.15 |
| NLys (K) | 0.314 | 8728.84 |
| NArg (R) | 0.311 | 5038.15 |
| NHis (H) | 0.305 | 4746.30 |
| NPhe (F) | 0.342 | 7415.82 |
| NTyr (Y) | 0.289 | 11655.32 |
| NTrp (W) | 0.297 | 9430.50 |

**Table S3.** Bonded Parameters for $BB^{-}$–BB–$BB^{+}$ Angles Across 18 Residues.

| **Residue Name** | **$BB^{-}$–BB–$BB^{+}$ Bong Angles (rad)** | **Force Constant (kJ/mol·$rad^2$)** |
|---|---|---|
| NAla (A) | 2.288 | 22.28 |
| NCys (C) | 2.298 | 20.33 |
| NAsp (D) | 2.311 | 24.21 |
| NGlu (E) | 2.302 | 22.28 |
| NIle (I) | 2.322 | 30.02 |
| NLeu (L) | 2.314 | 20.60 |
| NMet (M) | 2.296 | 25.31 |
| NAsn (N) | 2.299 | 20.22 |
| NGln (Q) | 2.302 | 23.69 |
| NSer (S) | 2.290 | 19.86 |
| NThr (T) | 2.347 | 25.23 |
| NVal (V) | 2.332 | 31.40 |

| | | |
|---|---|---|
| NLys (K) | 2.309 | 22.04 |
| NArg (R) | 2.330 | 23.59 |
| NHis (H) | 2.296 | 21.21 |
| NPhe (F) | 2.304 | 23.47 |
| NTyr (Y) | 2.299 | 21.52 |
| NTrp (W) | 2.323 | 25.29 |

**Table S4.** Bonded Parameters for BB⁻–BB–SC1 Angles Across 18 Residues.

| Residue Name | BB⁻–BB–SC1 Bong Angles (rad) | Force Constant (kJ/mol·rad²) |
|---|---|---|
| NAla (A) | 1.136 | 334.06 |
| NCys (C) | 1.327 | 111.90 |
| NAsp (D) | 1.172 | 340.27 |
| NGlu (E) | 1.313 | 193.10 |
| NIle (I) | 1.404 | 227.12 |
| NLeu (L) | 1.240 | 424.16 |
| NMet (M) | 1.381 | 89.62 |
| NAsn (N) | 1.279 | 211.20 |
| NGln (Q) | 1.318 | 125.43 |
| NSer (S) | 1.147 | 284.53 |
| NThr (T) | 1.309 | 328.00 |
| NVal (V) | 1.384 | 356.29 |
| NLys (K) | 1.275 | 326.50 |
| NArg (R) | 1.277 | 359.00 |
| NHis (H) | 1.149 | 363.95 |
| NPhe (F) | 1.209 | 135.53 |
| NTyr (Y) | 1.216 | 651.98 |
| NTrp (W) | 1.179 | 231.59 |

**Table S5.** Bonded Parameters for SC1 –BB–BB⁺ Angles Across 18 Residues.

| Residue Name | SC1 –BB–BB⁺ Bong Angles (rad) | Force Constant (kJ/mol·rad²) |
|---|---|---|

| | | |
|---|---|---|
| NAla (A) | 2.694 | 11.39 |
| NCys (C) | 2.366 | 15.34 |
| NAsp (D) | 2.523 | 21.15 |
| NGlu (E) | 2.436 | 21.54 |
| NIle (I) | 2.304 | 26.99 |
| NLeu (L) | 2.422 | 20.81 |
| NMet (M) | 2.348 | 18.42 |
| NAsn (N) | 2.500 | 17.56 |
| NGln (Q) | 2.424 | 18.17 |
| NSer (S) | 2.548 | 12.45 |
| NThr (T) | 2.321 | 23.97 |
| NVal (V) | 2.307 | 27.80 |
| NLys (K) | 2.504 | 17.41 |
| NArg (R) | 2.448 | 18.51 |
| NHis (H) | 2.553 | 17.49 |
| NPhe (F) | 2.458 | 19.84 |
| NTyr (Y) | 2.553 | 15.32 |
| NTrp (W) | 2.533 | 15.58 |

**1.2 Dihedral Parameters for Each Residue**

The dihedral potentials follow an Ryckaert-Bellemans (RB) form rather than a simple harmonic one, making direct parameter comparison impractical. We therefore use pointwise averages and standard deviations to capture their variability, as shown in the following figure.

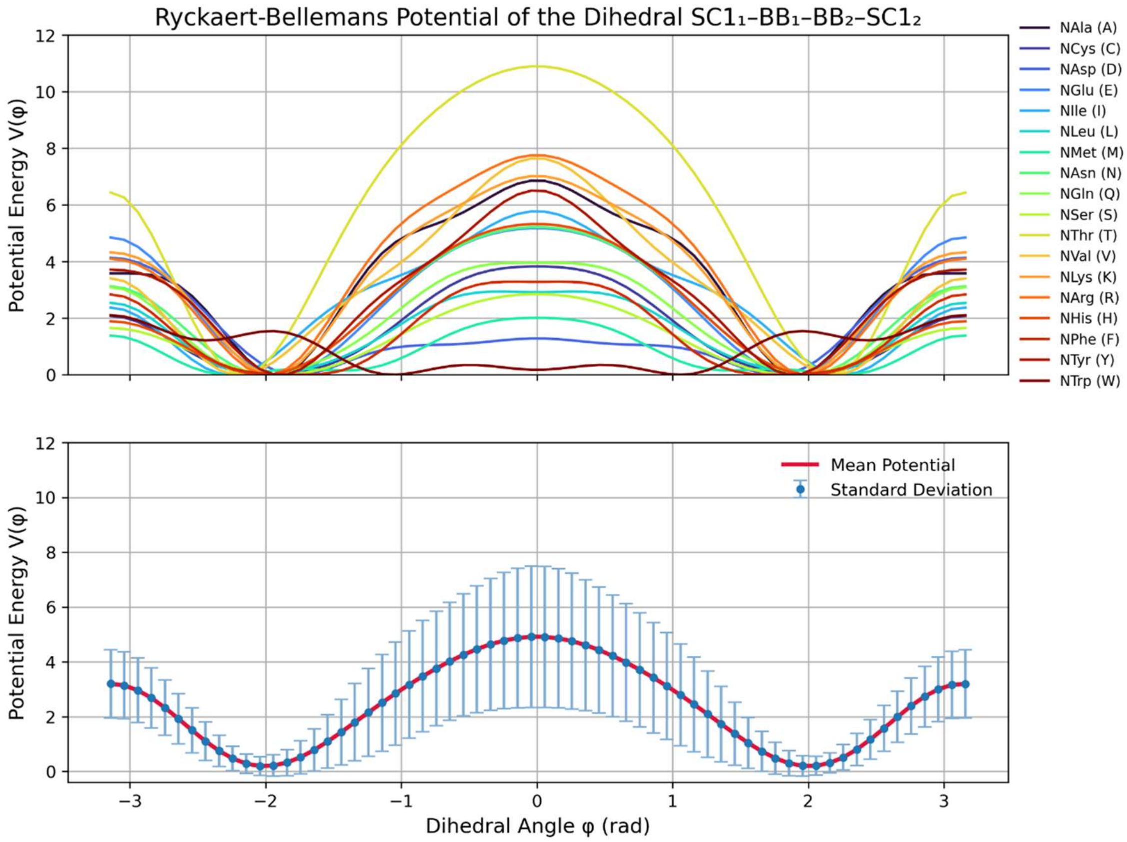


**Figure S1. Ryckaert-Bellemans Dihedral Potentials for the SC–BB–BB$^{+}$–SC$^{+}$ Backbone Torsions.** (Top panels) Residue specific Ryckaert-Bellemans dihedral potentials for the four backbone dihedral angles. Each colored curve corresponds to the fitted RB potential for one of the 18 peptoid residues. (Bottom panels) Averaged RB potentials for each dihedral angle, shown together with standard deviations. The averaged profiles exhibit consistent minima and overall shapes across residues, supporting the use of a unified dihedral representation for the CG model.

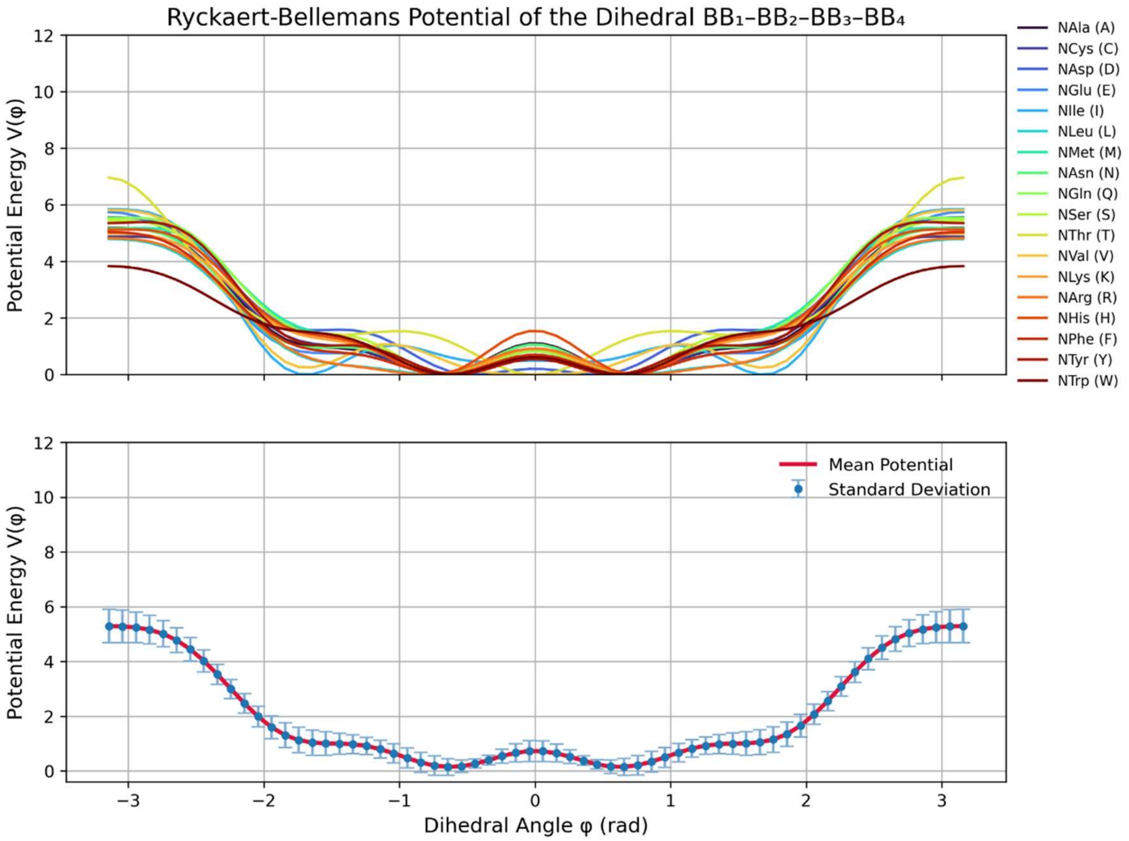


**Figure S2. Ryckaert–Bellemans Dihedral Potentials for the $BB^{-}$–BB–$BB^{+}$–$BB^{++}$ Backbone Torsions.** (Top panels) Residue specific Ryckaert-Bellemans dihedral potentials for the four backbone dihedral angles. Each colored curve corresponds to the fitted RB potential for one of the 18 peptoid residues. (Bottom panels) Averaged RB potentials for each dihedral angle, shown together with standard deviations. The averaged profiles exhibit consistent minima and overall shapes across residues, supporting the use of a unified dihedral representation for the CG model.

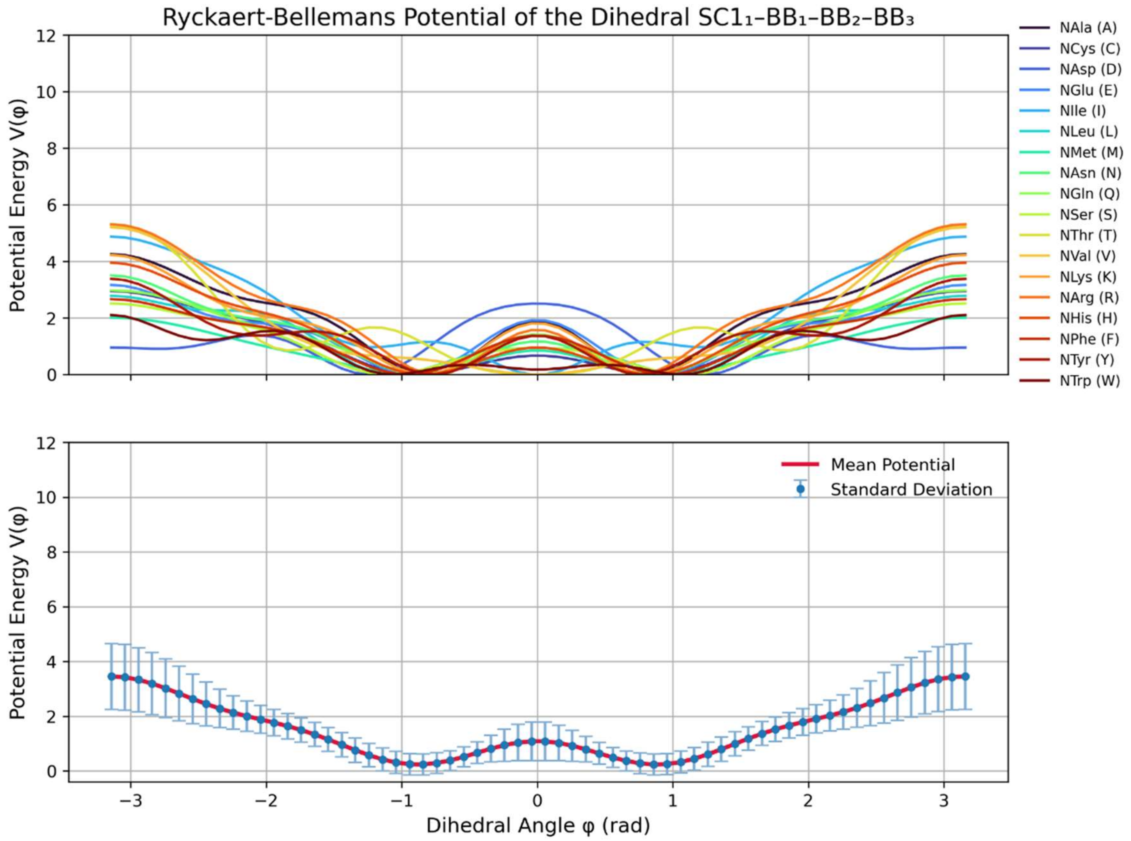


**Figure S3. Ryckaert–Bellemans Dihedral Potentials for the $SC^{-}$–$BB^{-}$–BB–$BB^{+}$ Backbone Torsions.** (Top panels) Residue specific Ryckaert-Bellemans dihedral potentials for the four backbone dihedral angles. Each colored curve corresponds to the fitted RB potential for one of the 18 peptoid residues. (Bottom panels) Averaged RB potentials for each dihedral angle, shown together with standard deviations. The averaged profiles exhibit consistent minima and overall shapes across residues, supporting the use of a unified dihedral representation for the CG model.

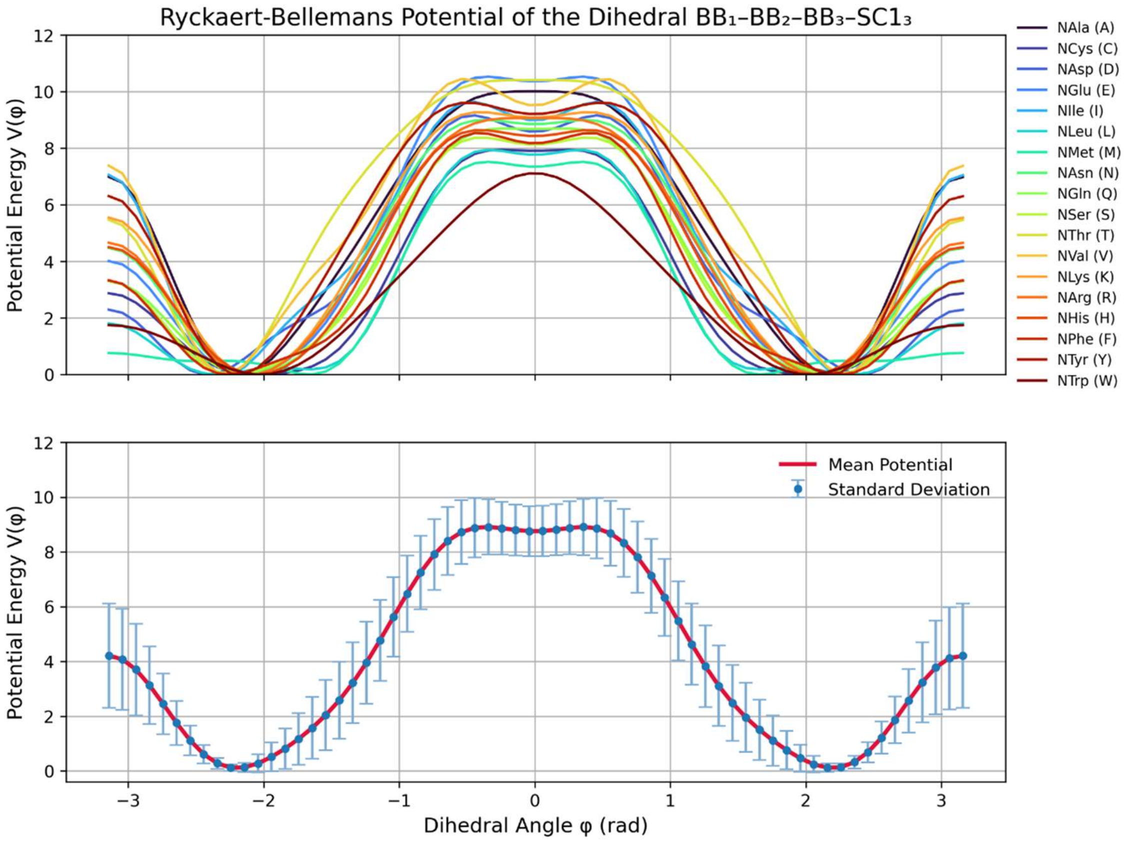


**Figure S4. Ryckaert–Bellemans Dihedral Potentials for the BB⁻–BB–BB⁺–SC⁺ Backbone Torsions.** (Top panels) Residue specific Ryckaert-Bellemans dihedral potentials for the four backbone dihedral angles. Each colored curve corresponds to the fitted RB potential for one of the 18 peptoid residues. (Bottom panels) Averaged RB potentials for each dihedral angle, shown together with standard deviations. The averaged profiles exhibit consistent minima and overall shapes across residues, supporting the use of a unified dihedral representation for the CG model.

### 2. Effects of Improper Dihedral Interactions

To further investigate the origin of the anomalously small Rg observed for NAsp, we found that it may be associated with improper dihedral $BB^{-}$–BB–SC1–$BB^{+}$ behavior. In particular, the distribution of this improper dihedral in the all-atom model deviates noticeably from that captured by the coarse-grained representation, leading to a more compact conformational preference. The specific features are as follows:

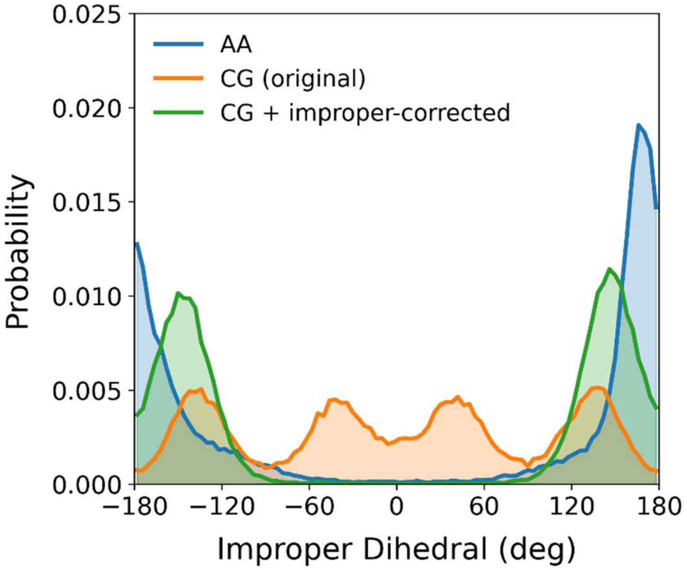


**Figure S5. Improper dihedral distributions for NAsp residues obtained from AA reference simulations and the corresponding CG model.** In the absence of explicit improper terms, the CG model fails to capture the main maxima and minima and exhibits a significant AA-CG discrepancy, likely due to the charged bead being directly connected to the backbone. This discrepancy is markedly improved upon introducing improper dihedrals.
